# Atheroprone Flow Activates SMAD-FOXO1 to drive Endothelial-to-Mesenchymal Transition and Atherosclerosis

**DOI:** 10.1101/2024.11.25.625323

**Authors:** Paul-Lennard Mendez, Lion Raaz, Jerome Jatzlau, Yunyun Xiao, Pieter Vrancaert, Jakub Ksiazkiewicz, Jakob Wallentin, Michael Trumpp, Stefanie Scharnitzki, Stefan Mundlos, Aernout Luttun, An Zwijsen, Fumiko Itoh, Petra Knaus

## Abstract

**Background:** Cardiovascular diseases are the leading cause of death worldwide with atherosclerosis as the main underlying pathology. A hallmark of atherosclerotic lesion formation is endothelial-to-mesenchymal transition (EndoMT) triggered by perturbed blood flow patterns at arterial bifurcations and curvatures. SMAD transcription factors (TFs), activated by bone morphogenetic protein (BMP) 9/10 or transforming growth factor beta (TGFβ) signaling, are indispensable for endothelial homeostasis. Yet, they also play a significant role in stimulating EndoMT. How different interacting co-factors mediate the shift towards a pathological SMAD response remains elusive.

**Methods:** We generated endothelial cell (EC)-specific SMAD1/5 or SMAD2/3 knock-out mice and performed assay for transposase accessible chromatin sequencing (ATAC-Seq) of EC nuclei from regions of atheroprone (aortic arch) and atheroprotective (descending thoracic aorta) flow to identify transcriptional co-regulators of SMADs. We validated this using single-cell (sc)ATAC-Seq and immunofluorescence staining data from wild-type mice. To assess conservation of our findings for the human situation, we performed co-immunoprecipitation and proximity ligation assays in human aortic ECs (HAoECs). We exposed HAoECs to pathological or physiological (*i*.*e*. oscillatory or pulsatile) flow and performed ATAC- and RNA-Seq. Next, transcriptomic and chromatin accessibility data were integrated and motif enrichment and TF footprinting analysis were performed. Finally, we used siRNA-mediated approaches, TF inhibition, and luciferase-based reporter gene assays to analyze the transcriptional response of target TFs and explored their presence in plaques of atheroprone low-density lipoprotein receptor-deficient mice.

**Results:** We observed enrichment of FOXO TF family motifs in DNA loci with increased accessibility in response to atheroprone flow *in vitro* and *in vivo*. These motifs were associated with genes displaying enhanced mRNA expression. We observed that FOXO motifs are enriched in peaks lost upon EC-specific SMAD KO in mice. We identified SMADs and FOXO1 as interacting partners that form complexes upon atheroprone flow stimulation. Inhibitor experiments revealed that FOXO1 and SMADs mediate EndoMT upon atheroprone flow exposure by upregulating *SNAI2*.

**Conclusion:** We identified SMAD/FOXO1 complexes that mediate EndoMT in response to atheroprone flow. Targeting this interaction can potentially reduce atherosclerotic burden by interfering with pathological flow-induced EndoMT and thus disease progression.

## Introduction

Cardiovascular diseases (CVDs) account for the majority of global deaths^1^ and atherosclerosis, the pathogenic build-up of cholesterol-rich plaques in larger arteries, is the main underlying cause^2^. Interestingly, atherosclerosis develops primarily at vasculature sites exposed to irregular hemodynamics, commonly referred to as **disturbed flow** (**DF**), while areas experiencing laminar flow (**LF**) are relatively protected from disease^3, 4^. As modeling DF *in vitro* is challenging, low, **oscillatory flow** (**OS**) is widely used to mimic this condition using **pulsatile flow** (**PS**) as a control. In both, *in vitro* and *in vivo* studies, pathogenic effects of irregular hemodynamics have been attributed to dysregulation of growth factor signaling pathways, including the bone/body morphogenetic protein (BMP) and transforming growth factor beta (TGFβ) family^5-8^.

BMP/TGFβ signaling mediates both, atheroprotective and atheroprone mechano-responses^7-10^. In particular, BMP/TGFβ signaling is involved in tissue homeostasis but also the trans-differentiation of endothelial cells (ECs) to mesenchymal cells in a process termed endothelial-to-mesenchymal transition (EndoMT), which underlies the onset and progression of atherosclerotic disease^11, 12^. As BMP and TGFβ ligands all signal through activation of a common limited set of SMAD transcription factors (TFs) – predominantly SMAD1/5 for BMPs, SMAD2/3 for TGFβ - it is intriguing that they induce such opposing processes. Thus, it has long been suggested that the outcome of BMP/TGFβ signaling is mediated by cell and tissue context-dependent co-factors that bind to SMAD TFs^13^. Which SMAD co-factors are responsible for directing the atheroprone BMP/TGFβ response is currently unknown.

Forkhead Box O (FOXO) TFs are important regulators of cell survival and proliferation^14^. Amongst the vascular FOXO members, FOXO1 has been described as a modulator of atherosclerotic disease^15-19^. FOXO members are phosphorylated and thereby inactivated downstream of PI3K/ AKT signaling, which mediates cytoplasmic retention^20^. FOXO1/3/4 have been shown to act as SMAD co-factors through interaction with SMAD3 and SMAD4 in human keratinocytes^21^. Importantly, FOXO proteins have pioneering functions, allowing them to bind to closed chromatin regions, to perturb core histones, and thus to make DNA accessible for other DNA binding factors^22^.

In this study, we demonstrate that SMADs and FOXO1 interact in response to atheroprone flow, collaboratively driving the expression of genes associated with atheroprone EndoMT through the regulation of *SNAI2*. Additionally, our findings indicate that the transcriptional activity induced by SMADs can either be inhibited or enhanced, depending on the activation state of FOXO1. These insights illuminate the intricate gene regulatory mechanisms underlying atheroprone flow-induced transcription and may provide a novel therapeutic option for treatment of atherosclerosis by targeting SMAD-FOXO1 interactions.

## Methods

### SMAD knock-out mice

SMAD1^flox/flox^; SMAD5^flox/flox^ (SMAD1/5^WT^), SMAD2^flox/flox^; SMAD3^flox/flox^ (SMAD2/3^WT^), and PDGFB-iCreER mice were maintained on a C57BL/6J background. Tamoxifen (Tx) (T5648; Sigma-Aldrich) dissolved in corn oil (20 mg/mL), was intraperitoneally administered (40 mg/kg/day) for 3 consecutive days into control (SMAD1/5^WT^ or SMAD2/3^WT^) and PDGFB-iCreER SMAD1/5^flox/flox^ or SMAD2/3^flox/flox^ mice, the latter resulted in EC-specific knockout (KO) mice of SMAD1/5 or SMAD2/3, named respectively SMAD1/5^iΔEC^ or SMAD2/3^iΔEC^. All the animal experiments were approved by the animal study committee (Tokyo University of Pharmacy and Life Sciences) and performed in accordance with the Guidelines of the Tokyo University of Pharmacy and Life Sciences for Animal and Recombinant DNA Experiments (L23-15). The mice were housed in the animal facilities of Tokyo University of Pharmacy and Life Sciences under specific pathogen-free conditions with constant temperature and humidity and were provided a standard diet. For experiments, adult male mice weighing 24–30 g were used.

To isolate EC sheets, the aortic arch and descending thoracic aorta were rapidly excised from the mouse and any adhering fat and connective tissue was carefully removed. They were then cut into 5 mm rings. Each aortic ring was opened by making an incision on the side opposite to the target area, allowing the intimal sheet to be peeled off under a stereomicroscope. The desired section was then trimmed to approximately 3 mm^2^ and immediately stored in liquid nitrogen for subsequent analysis.

### LDLR Knock-out and WT C57BL/6 mice

Low-density lipoprotein receptor (LDLR) knock-out mice^23^ and their wild-type (WT) littermates were housed under standard conditions in IVC-cages with controlled temperature of 20 °C ± 3 °C. The mice were kept in a 12/12-h light/dark cycle and allowed access to standard rodent chow and water ad libitum. Before the dissections, the mice were perfused with a saline solution. The dissected tissues were then snap-frozen in liquid N_2_ and stored at -80 °C until cryosectioning. Samples were mounted with Tissue-Tek (Sakura), and 7-μm sections were made using a Leica 3050S cryostat. All animal experiments were approved by the KU Leuven ethical committee for animal experimentation under ECD number CVW-061/2021.

### Immunofluorescence staining of mouse cryosections

Cryosections were equilibrated to room temperature for 60 min and rehydrated in PBS for 5 min. Subsequently, sections were fixed in 4 % Paraformaldehyde (PFA) in PBS for 10 min at 4 °C. Upon washing twice with PBS for 5 min, tissue sections were blocked for 20 min with 10 % normal goat serum in PBS containing 0.3 % Triton-X-100 (blocking buffer). Primary antibodies (anti-SMAD1 XP, Cell Signaling Technology 6944, 1:100; anti-FOXO1, Cell Signaling Technology 2880, 1:100) were added in blocking buffer and incubated overnight at 4 °C. Control slides were incubated in blocking buffer without primary antibodies. Sections were then washed 3 times with 0.3 % Triton-X-100 in PBS, incubated with secondary antibody (goat anti-rabbit Alexa Fluor 647, Invitrogen A21244, 1:300) for 1 h at room temperature, followed by 3 washes in PBS-Triton-X-100. Finally, sections were incubated with DAPI in PBS (D9542, Sigma-Aldrich, 1:500) for 5 min, washed once in PBS-Triton-X-100 for 5 min and mounted using ProLong Diamond anti-fade (Invitrogen). Images were captured on a Leica TCS SP8 confocal microscope equipped with a 405-nm diode laser and 633nm HeNe laser using a 40× immersion oil objective.

### Single-Cell ATAC-Seq

FASTQ files were downloaded via the SRA database (Accession number PRJNA646233) and aligned to the murine genome (Cell Ranger mm10-2020-A-2.0.0) using cellranger-atac -count command (10x Genomics Cell Ranger ATAC v2.0.0). Cell Ranger outputs were loaded into R using Signac^24^ and Seurat^25^. Gene activity scores were calculated and used as measures of gene expression to identify EC clusters based on expression of *Cdh5*, *Pecam1*, *Icam2*, and *Tie1*. EC clusters were subsetted and motif activities of Smad and FOXO were calculated using ChromVar^26^. Visualization of motif activities was performed using ggplot2^27^.

### Cell culture

Human Aortic Endothelial Cells (HAoECs; Cell Applications, USA) from three different donors (1576, 2939, 2986) were cultured in Endothelial Cell Growth Medium 2 (EGM2, C-22111, PromoCell GmbH, Germany) supplemented with 2 % fetal calf serum (FCS), and penicillin (100 units/ mL) / streptomycin (100 µg/ mL) in a humidified atmosphere at 37 °C and 5 % CO_2_ (v/v). Cells were cultured on 0.1 % (w/ v) pork skin gelatine (Sigma Aldrich) in PBS for general cell culture and experiments. Cells were analyzed between passages three and five. Human THP-1 monocytes were cultured in RPMI 1640 medium (Gibco; ThermoFisher Scientific, Inc., MA, USA) containing 10 % FCS, 2 mM l-glutamine, 100 U/ mL penicillin, and 100 μg/ mL streptomycin (Gibco; ThermoFisher Scientific, MA, USA).

### Transient transfection with siRNA

siRNAs were purchased from Dharmacon (ON-TARGET plus Non-targeting siRNA #1, D-001810-01-50; ON-TARGET plus Human FOXO1; J-003006-09-0002). HAoECs were transfected using Lipofectamine2000 (ThermoFisher Scientifc) with 40 nM siRNA in 6-well plates (400.000 cells/ well, 2 mL medium) according to the manufacturer’s instructions. Twenty-four hours (h) after transfection, cells were trypsinized and reseeded to μ-Slides I 0.4 (ibidi) at 3.5*10^6^ cells/ mL. The day after reseeding, FSS (24 h) experiments were started. Thus, cells were harvested 3 days after siRNA transfection.

### Real-time quantitative PCR (RT-qPCR)

Cellular RNA was isolated using the NucleoSpin RNA XS isolation kit (Macherey-Nagel) according to the manufacturer’s instructions. Total RNA was reversely transcribed by incubating it with random primers (100 pmol μL^-1^, Invitrogen) and M-MuLV reverse transcriptase enzyme (200,000 U mL^-1^, New England Biolabs) was added per sample. RT-qPCR was performed using a StepOnePlus Real-Time PCR System (ThermoFisher Scientific) with specific primers listed below. Reactions were performed in triplicates in MicroAmp Optical 96-well reaction plates (ThermoFisher Scientific) using Luna PCR Master Mix (New England Biolabs). Fold induction was calculated by comparing relative gene expression to the housekeeping gene *RSP9* using the ΔΔCT method.

### Application of fluid shear stress

First, 3.5*10^5^ cells were seeded in EGM-2 to ibidi µ-Slide I Luer 0.4 mm (ibidi GmbH, no. 80176) pre-coated with 0.1 % pork skin gelatin (Sigma-Aldrich). Next, cells were kept under culture conditions (37°C, 5 % CO_2_, 95 % RH) for 48 h with daily medium exchanges. Oscillatory FSS (OS; ± 5 dyn/ cm^2^, 1 Hz, changing directions) or pulsatile laminar FSS (PS; 20 dyn/ cm^2^, 1 Hz, unidirectional) were applied for 24 h using a pneumatic pump system (ibidi GmbH, no. 10902) with the associated software (v.1.6.0).

### Proximity Ligation Assay (PLA)

PLA on flow-exposed HAoECs was performed as previously described^28^. In short, HAoECs were subjected to OS or PS (see above) for 24 h, washed with PBS, and fixed in 4 % PFA in PBS. Subsequently, cells were permeabilized with 0.3 % Triton-X-100 in PBS for 10 min and blocked with Duolink blocking solution (Sigma Aldrich). Antibodies used were anti-FOXO1 (Cell Signaling, C29H4, Rabbit mAb #2880; dilution 1:100) and anti-SMAD1 (abcam, 913C1b, ab53745, dilution 1:100). Samples were then incubated with (+)-mouse and (-)-rabbit secondary probes and amplification was performed with Duolink Amplification buffer Red (Sigma Aldrich). Cells were counterstained with DAPI, and images were captured on a Leica TCS SP8 confocal microscope equipped with a 405-nm diode laser, 488-nm argon laser, 561-nm diode laser, and 633nm HeNe laser using a 20× (20×/0.75 HC PL APO Imm Corr WD 0.68 mm) immersion oil objective. Image analysis was performed with FIJI using the count particles function^29^.

### Reporter Gene Assays

To generate the firefly luciferase reporters, forward and reverse oligonucleotides were annealed using 1x T4 Ligation buffer at 95 °C and then phosphorylated with a T4 polynucleotide kinase. The annealed and phosphorylated oligos were then ligated using T4 ligase into a pGL3-basic minimal adenoviral major late promoter (MLP) vector using *KpnI* and *XhoI* restriction sites upstream of the MLP. The ligation was heat inactivated at 65 °C for 10 min followed by transformation using chemically competent DH5α *E. coli* bacteria. Plasmid DNA purification was carried out using NucleoSpin Plasmid EasyPure Kit following the manufacturer’s guidelines, followed by construct validation by Sanger sequencing. HEK293T cells were transfected using Polyethylenimine (PEI) with respective luciferase reporter constructs. A constitutively expressing construct encoding renilla luciferase (RL-TK; Promega) was co-transfected as an internal control and an empty vector or a vector containing FOXO1A-FLAG^30^. On the following day, cells were starved in serum-free medium for 3 h and then stimulated with 5 nM BMP6 for 24 h. Cells were lysed using passive lysis buffer (Promega) and luciferase activity was measured with a TECAN Spark Luminometer, following the manufacturer’s instructions. FOXO1A-FLAG was a gift from Stefan Koch (Addgene plasmid # 153141; http://n2t.net/addgene:153141; RRID:Addgene_153141).

### Co-immunoprecipitation

Co-immunoprecipitation was performed as described elsewhere^31^. Cells were lysed in RIPA buffer (150 mM NaCl, 50 mM Tris/HCl pH 7.4, 0.1 % SDS, 0.5–1% NP-40/ IGEPAL) supplemented with protease/phosphatase inhibitors (1 mM PMSF, 2 mM Na_3_VO_4_, 20 mM Na_4_P_2_O_7_, 50 mM NaF, complete protease inhibitor cocktail (Roche)). Immunoprecipitation was performed using either 3 µg of either SMAD1 (SMAD1 XP, Cell Signaling Technology 6944), SMAD2/3 (SMAD 2/3 XP, Cell Signaling Technology 8685) or isotype control (Rabbit IgG XP, Cell Signaling Technology 3900) antibody overnight followed by incubation with protein A coupled Sepharose beads (VWR). Samples were washed with lysis buffer, eluted with Laemmli buffer, and subjected to Western blotting.

### Western Blotting

Samples in Laemmli buffer^32^ were denatured at 95 °C for 10 min before separation on 10 % SDS–PAGE gels. Proteins were transferred onto PVDF membranes (Roth). Membranes were blocked in 3 % w/ v BSA (Sigma-Aldrich) in TBST and then incubated with primary antibodies overnight at 4 °C. The next day, membranes were incubated with goat-α-rabbit-HRP (1:5,000, Dianova, Hamburg, Germany), before detection with WesternBright Quantum ECL HRP substrate (advansta) using a Fusion-FX7 (Vilber Lourmat). Primary antibodies used were against FOXO1 (Cell Signaling Technology, C29H4, Rabbit mAb #2880), SMAD1 (3267, Cell Signaling Technology), SMAD2/3 (Cell Signaling, D7G7, Rabbit mAb #8685), and GAPDH (Cell Signaling Technology 2118). The concentration of primary antibodies was 1:1,000 in 3 % BSA.

### Monocyte adhesion assay

Monocyte adhesion assay was performed as previously described^7^. In brief, HAoECs were subjected to OS and PS for 24 h. Before the end of the FSS experiment, THP-1 cells per were stained with 5 µM cell tracker green (Invitrogen, C2925) in RPMI 1640 medium without additives for 30 min. Subsequently, cells were washed 3 times with PBS and resuspended at 3* 10^6^ cells/ mL in EGM2 medium. After the end of FSS experiments, µ-slides containing HAoECs were detached from the flow setup, excess medium from the reservoirs of µ-slides was aspirated and 300 µL stained THP-1 cell suspension was added in one reservoir, while aspirating from the other reservoir, efficiently leaving 100 µL of cell suspension in the channel. THP-1 cells were incubated on HAoEC monolayers for 30 min, followed by a gentle wash step with PBS and fixation in 4 % PFA in PBS for 10 min. For each experiment and condition, 5 images were acquired with a fluorescence phase-contrast microscope (Zeiss Axio Observer). Adhered leukocytes were counted with FIJI software and data of each donor was normalized to the overall mean of PS.

### ATAC-Seq on human aortic endothelial cells

ATAC-Seq of FSS exposed cells was performed as described before^9, 33^. In brief, 2 µ-Slides I Luer 0.4 mm per donor and condition were cut open with a scalpel, and adherent cells were trypsinized and collected by centrifugation in EBM2 medium. For each sample, 50,000 cells were transposed and cleaved DNA was purified using the MinElute Reaction Cleanup Kit (Qiagen, no. 28206). For clean-up, the manufacturer’s protocol was followed with two exceptions. First, elution was done in 11 µL to get a full 10 µL residual volume, and second, a longer elution time (3-5 min) was used for a higher yield of DNA fragments. DNA concentration was measured with a Qubit fluorimeter. DNA fragments were stored at -20°C until further processing. For FOXO1 inhibition ATAC-Seq, 0.5 µM AS1842856 in DMSO was added to the cell culture medium 1 h prior to flow stimulation. Similarly, DMSO was added to control cells 1 h prior to flow.

### ATAC-Seq on mouse tissue

ATAC-Seq of mouse tissue was performed according to a protocol modified from ref.^34^. Tissue was placed in 1 mL ice-cold homogenization buffer (0.26 M Sucrose, 0.03 M KCl, 0.001 M MgCl_2_, 0.02 M Tricine-KOH pH 7.8, 0.001 M DTT, 0.5mM Spermidine, 0.15 mM Spermine, 0.3 % IGEPAL/ NP40, cOmplete Protease Inhibitor Tablet, ad H_2_O) in a chilled 2 mL douncer (Kimble), left to thaw for 5 min and dounced with pestle A until resistance was gone (approximately 10 strokes). The tissue was then dounced with Pestle B for 20 strokes. Cell suspensions were then divided into two equal parts, collected in pre-chilled 1.5 mL reaction tubes (Eppendorf, Germany) and centrifuged at 4 °C for 10 min at 500 rcf. Supernatant was carefully removed, cells were resuspended in 50 µL ATAC Reaction mix (1 µL Tn5 (Illumina), 6 µL H_2_O, 16.5 µL PBS, 25 µL 2xTD buffer (Illumina), 0.5 µL 10 % Tween20, 0.5 µL 1 % Digitonin) and incubated at 37 °C with 1,000 rpm constant shaking for 30 min. DNA was purified using the MinElute Reaction Cleanup Kit (Qiagen, no. 28206) as described above.

### ATAC-seq library preparation

Library generation, amplification and purification was conducted as in Buenrostro *et al*.^35^, including the RT-qPCR step to estimate the appropriate number of additional PCR cycles. In brief, indexing primers v2_Ad1.2/3 and v2_Ad2.3/4/5/6 (2.5 µL each) from Buenrostro *et al*.^36^ were ligated to the DNA fragments (10 µL + 10 µL H_2_O) using PCR Master Mix (NEB, M0541L) (25 µL) in a thermocycler (72 °C, 5 min; 98 °C, 30 s; 5x (98 °C, 10 s; 63 °C, 30 s; 72 °C 1 min)). Quantitative PCR was conducted to determine the final number of PCR cycles with 5 µL Library, 2.5 µL H_2_O, 0.5 µL Ad1.x, 0.5 µL Ad2.x, 1.5 µL 10x SYBR Green I (ThermoFisher Scientific, S7563), 5 µL PCR Master Mix with the above-mentioned settings for 20 cycles, skipping the initial 72°C, 5 min. The partially amplified libraries were then further cycled with the above-mentioned settings, according to the CT values from the qPCR. After that, two-sided size selection with magnetic AMPure XP Beads (Beckmann Coulter, no. A63881; 0.55× and 0.9× sample volume of bead solution added) was used to remove primer-dimers and large (> 1 kb) DNA fragments. Library quality was assessed with an HS DNA Bioanalyzer chip (Agilent, no. 5067-4626) before 2×100 paired-end Illumina high output sequencing (Max Planck Sequencing Core Facility at MPIMG).

### ATAC-seq data preprocessing

ATAC-seq data were processed via the standard ENCODE ATAC-seq pipeline, using Caper with Conda (v1.10.0, https://github.com/ENCODE-DCC/atac-seq-pipeline/releases/tag/v1.10.0). Briefly, reads were aligned with Bowtie2 (v2.3.4.3)^37^ to the hg38 reference genome and filtered for unmapped, duplicates and mitochondrial reads. Peak calling was performed using MACS2 (v2.2.4)^38^ for each replicate.

### ATAC-seq data analysis

To determine differentially accessible peaks in mouse and HAoEC FOXO1 inhibition ATAC-Seq data we used csaw^39^. We first defined a consensus peak set of all conditions and then counted reads in 300bp windows. Subsequently, counts were Loess normalized and differentially accessible regions (DARs) were determined with edgeR^40^. To determine differentially accessible peaks in HAoEC OS *versus* PS ATAC-Seq data we constructed read count matrices of consensus peaks using featureCounts (v2.0.1) and normalized counts using RUVr (k=3) from the RUVseq package (v1.28.0)^41^ according to Gontarz *et al.*^42^. Subsequently, differentially accessible regions (fold change > 1.5, adjusted p-value < 0.05) were determined with DeSeq2 (v1.34.0)^43^. Gene Ontology analysis of DARs was performed with GREAT tool (v4.0.4)^44^ in basal plus extension mode with default settings. ATAC-Seq tracks were visualized with sparK^45^. To analyze motif occurrences, we used *annotatepeaks* module with -find and built-in hg38 genome from HOMER^46^ for FOXO1 and SMAD motif (*GTCTG*) on OS DARs (fold change > 1.5, adjusted p-value < 0.05). Motif enrichment analysis was performed using HOMER’s *findmotifsgenome* script with -size given -mask flags on differentially accessible regions.

### TF footprinting analysis

We employed TOBIAS (v0.15.0)^47^ framework using the TOBIAS snakemake pipeline to conduct the TF footprinting analysis with the HOMER TF database and added custom SMAD motifs (*GGTGCC*, *GGCGCC*, *GTCTG*). To perform TF footprinting, a config file with standard settings, referencing the GRCh38 assembly (GCF_000001405.26), blacklisted regions (ENCFF356LFX), and a consensus peak set, was created and the analysis was invoked on a linux cluster with snakemake --configfile TOBIAS_configuration.yaml --use-conda --cores 32.

### Cleavage Under Targets and Release Using Nuclease (CUT&RUN)

CUT&RUN was performed using a commercially available kit (Cell Signaling Technology) by following the accompanying protocol. For this, HAoECs were cultured in 6-well plates, starved for 3 h in EBM2 without FCS (PromoCell) and stimulated with 0.3 nM BMP9 for 90 min. Cells were washed with PBS, trypsinized and counted. Next, 5*10^5^ cells were permeabilized, Concavalin A beads added, and incubated with SMAD1 or SMAD2/3 antibody overnight at 4°C. The next day, DNA was digested using pAG-MNase for 30 min at 4°C. DNA was purified using spin columns (Cell Signaling Technology). Libraries were constructed using the NEBNext Ultra II DNA Library Prep Kit for Illumina (New England Biolabs) following a published protocol^48^. Final libraries were sequenced on an AVITI Sequencer (Element Biosciences) in PE75 mode. The resulting FASTQ files were processed using the standard ENCODE ChIP-Seq pipeline in histone mode, using Caper with Conda (https://github.com/ENCODE-DCC/chip-seq-pipeline2).

### RNA-Seq set-up

For RNA-Seq analysis, cells were simultaneously harvested with ATAC-Seq samples. One µ-Slide I Luer 0.4 mm per donor and condition was washed with PBS once and RNA was isolated as described here for RT-qPCR. After initial quality control using Agilent’s Bioanalyzer, sequencing libraries were prepared from 500 ng of total RNA per sample following Roche’s stranded “KAPA RNA HyperPrep” library preparation protocol for single indexed Illumina libraries. First, the polyA-RNA fraction was enriched using oligo-dT-probed paramagnetic beads. Enriched RNA was heat-fragmented and subjected to first-strand synthesis using random priming. The second strand was synthesized incorporating *dUTP* instead of *dTTP* to preserve strand information. Afterwards, A-tailing Illumina sequencing-compatible adapters were ligated. Following bead-based clean-up steps, the libraries were amplified using 10 cycles of PCR. Library quality and size were checked with Qubit, Agilent Bioanalyzer, and qPCR. Sequencing was carried out in biological triplicates on an Illumina NovaSeq 6000 system in PE101 (10|10) mode (paired end, 100 bp read length) yielding between 31 and 62 million fragments per sample.

### RNA-Seq data analysis

After initial quality control with FastQC^49^, RNA-Seq reads were aligned to the human genome (Gencode GRCh38.p13) and quantified with Salmon (v1.9.0)^50^. Differentially expressed genes (DEGs) were quantified with DESeq2 (v1.34.0)^43^. Data visualization was performed with RStudio^51^ environment for R (v4.1.1)^52^ using the following packages: ggplot2^27^, dplyr^53^, and ggrepel^54^.

### Integration of RNA-Seq and ATAC-Seq data

Highest overall peaks (|fold change| > 1.5, adjusted p-value < 0.05) of ATAC-Seq data were correlated with corresponding to DEGs (|fold change| > 1.5, adjusted p-value < 0.05). Motif enrichment analysis of the underlying peaks was performed using HOMER (v4.11.1) and *findMotifsGenome* module with - size 200 and -mask flags and built-in hg38 genome for *de novo* motif discovery.

### Figures

Schemes were created with BioRender.com and figures were assembled using Adobe Photoshop 2020.

### Statistical analysis

Statistical analyses were conducted using GraphPad Prism (v9.3). Specific tests for each analysis are provided in the corresponding figure legends. The Shapiro-Wilk test was applied to assess the normality of datasets. If normality could not be assumed, the Mann-Whitney U-test was used. For analyses involving normalization to a single condition, a one-sample t-test was conducted with the theoretical mean set to 1. Group comparisons across conditions were analyzed with a two-way ANOVA followed by Šídák’s post-hoc test. For RNA-Seq data, adjusted p-values calculated by DESeq2 were used for significance testing of normalized counts (Wald test with Bonferroni correction for multiple testing).

### Code and data availability

All relevant scripts for scATAC-Seq, ATAC-Seq, and RNA-Seq data analysis and visualization as well as PLA quantifications are accessible from https://github.com/pmadnez/SMAD-FOXO1. RNA-Seq, ATAC-Seq, and CUT&RUN-Seq raw and processed data are available from GEOXXXXXX.

## Results

### FOX motifs are enriched in SMAD-dependent chromatin accessibility peaks in mice

To investigate which co-factor(s) orchestrate(s) the atheroprone transcriptional response of SMADs, we generated EC-specific SMAD1/5 (*SMAD1/5^iΔEC^*) and SMAD2/3 (*SMAD2/3^iΔEC^*) knock-out (KO) mice by crossing *SMAD1/5^flox/flox^*(*SMAD1/5^W^*^T^ ref.^55, 56^) and *SMAD2/3^flox/flox^* (S*MAD2/3^WT^*ref.^57, 58^) mice with *PDGFB-iCreER*; *SMAD1/5^flox/flox^* or *PDGFB-iCreER*; *SMAD2/3^flox/flox^* (ref.^59^) mice (**Figure 1A, S1A**). We first analyzed chromatin accessibility in ECs isolated from *SMAD1/5^WT^* control mice from either the inner curvature of the aortic arch (DF) or the descending thoracic aorta (unidirectional laminar flow, LF) using bulk ATAC-Seq (**Figure 1B**). To minimize the mechanical stress during the harvesting process, we peeled off EC sheets at the indicated areas of fresh mouse aortae and immediately snap-froze them. In accordance with published scATAC-Seq data, we observed higher chromatin accessibility at the *Ccl2* and *Acta2* promoters in the DF samples while the promoter region of the laminar shear stress-sensitive gene *Cyp1b1* (ref.^60^) showed decreased accessibility, together proving the validity of our sampling approach (**Figure 1C**)^61^. Furthermore, we found the promoter region of BMP target gene *Id1* and an intronic region near the *Snai2* promoter, a well-known EndoMT-mediating TF^62^, to display higher accessibility in aortic arch ECs (**Figure 1D**). Intriguingly, accessibility of the *Id3* promoter did not display differences (**Figure 1D**). To understand how SMAD TFs regulate the endothelial response to DF, we next compared chromatin accessibility in aortic arch ECs of *SMAD1/5^WT^* and *SMAD1/5^iΔEC^* mice via ATAC-Seq (**Figure 1E**). First, regions that were no longer accessible upon EC-specific KO of SMAD1/5 were highly enriched for established SMAD motifs (**Figure 1F**). Second, chromatin accessibility at the intronic region of *Snai2* identified before and the *Id1* promoter, a known target of SMAD1/5 TFs in ECs^63^, was decreased upon SMAD1/5 KO in *SMAD1/5^iΔEC^* mice (**Figure 1G**). Interestingly, chromatin accessibility at the *Id3* locus was enhanced by SMAD1/5 KO, showcasing the complex, likely co-factor-dependent regulation by SMAD TFs (**Figure 1G**). Finally, we performed motif enrichment analysis in peaks that were lost upon SMAD1/5 KO and screened for motifs of TFs known to physically interact with SMADs. Along with enrichment of SMAD-like motifs^9^, we observed the enrichment of FOX motifs (**Figure 1H**)^64-66^.

**Figure 1.**
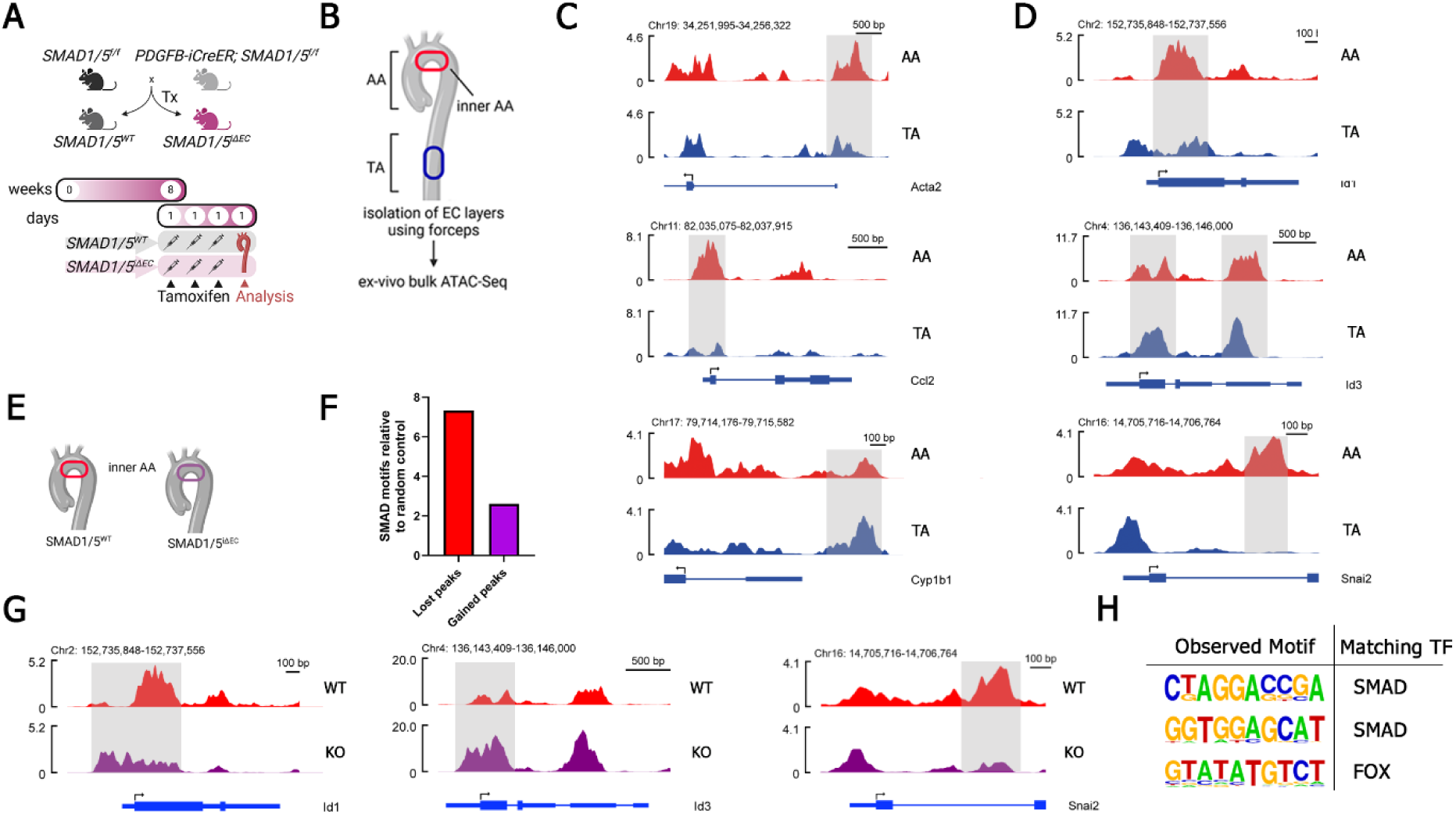
FOX motifs are enriched in SMAD1/5-dependent peaks in mouse endothelial cells (ECs). **A** Scheme of mouse breeding and tamoxifen injection. **B** Isolation of ECs from the inner curve of the aortic arch (AA, disturbed, multidirectional flow (DF)) or the descending thoracic aorta (TA, laminar, unidirectional (LF)) for bulk ATAC-Seq. **C,D** ATAC-Seq tracks of ECs isolated either from the TA (blue, LF) or AA (red, DF). **E** Isolation of ECs from the inner curvature of AA of *SMAD1/5^WT^* (red) or *SMAD1/5^iΔEC^* (pink) mice for ATAC-Seq (n= 2/3 *SMAD1/5^WT^* and *SMAD1/5^iΔEC^* mice). Gray shading highlights area of interest. **F** Number of SMAD motifs (*GGCGCC*) found in peaks differentially accessible peaks (FDR<0.2) compared to random genomic regions of similar size and number. **G** Genome browser view showing ATAC-Seq tracks of ECs isolated from the inner curvature of AA of WT (red) or SMAD1/5 knock-out (pink) mice. Gray shading highlights area of interest. **H** Motifs enriched in ATAC-Seq peaks that are lost upon SMAD1/5 knock-out identified by HOMER. Found motifs were screened for similarities to motifs of SMADs or known SMAD co-factors.

Next, we investigated chromatin accessibility profiles upon endothelial SMAD2/3 depletion in *SMAD2/3^iΔEC^* mice (**Figure S1A,B**). Again, we observed a strong enrichment of SMAD motifs in peaks lost upon SMAD2/3 KO (**Figure S1C**). Accessibility was decreased at the promoter of *Id1* and the promoter and the previously identified intronic region of *Snai2* (**Figure S1D**). In contrast, the *Id3* promoter region did not display decreased accessibility, similar to what we observed in *SMAD1/5^iΔEC mice^* (**Figure S1D**). Motif enrichment analysis of peaks lost upon SMAD2/3 KO again showed enrichment of FOX motifs along with SMAD motifs (**Figure S1E**).

In summary, we showed that both SMAD1/5 and SMAD2/3 regulate chromatin accessibility of *Snai2* in ECs of mice in response to atheroprone flow and identified FOX TFs as potential co-factors governing the atheroprone SMAD signaling.

### SMADs and FOXOs are active in ECs exposed to DF *in vivo*

To validate our findings from bulk ATAC-Seq experiments, we made use of existing scATAC-Seq data from a partial carotid artery ligation mouse model (**Figure 2A**)^61^. The data includes cells sampled 2 days (2d) or 2 weeks (2w) after partial ligation. Here, the right carotid artery serves as a unidirectional laminar flow control, while cells from the left carotid artery are exposed to DF. We performed cell clustering, assigned cell types through gene activities of cell type markers, and identified 7 distinct EC clusters (**Figure 2B**). Amongst those clusters, one was specific for chronic (2w) exposure to DF (EC3) and two were specific for the acute (2d) DF condition (EC4, EC6) (**Figure 2C**). We then performed motif activity analysis using the ChromVAR package^26^. We computed activities for a SMAD motif (JASPAR 2022, MA0535.1) and a FOX motif (JASPAR 2022, MA0480.1). When displaying the activity by cell on our clustered ECs, we observed the highest activity for both SMAD and FOX motifs in cluster EC3 which is mostly present in chronic DF, highlighting the co-activity of SMADs and FOX TFs in DF-exposed ECs *in vivo* (**Figure 2D,E**).

**Figure 2.**
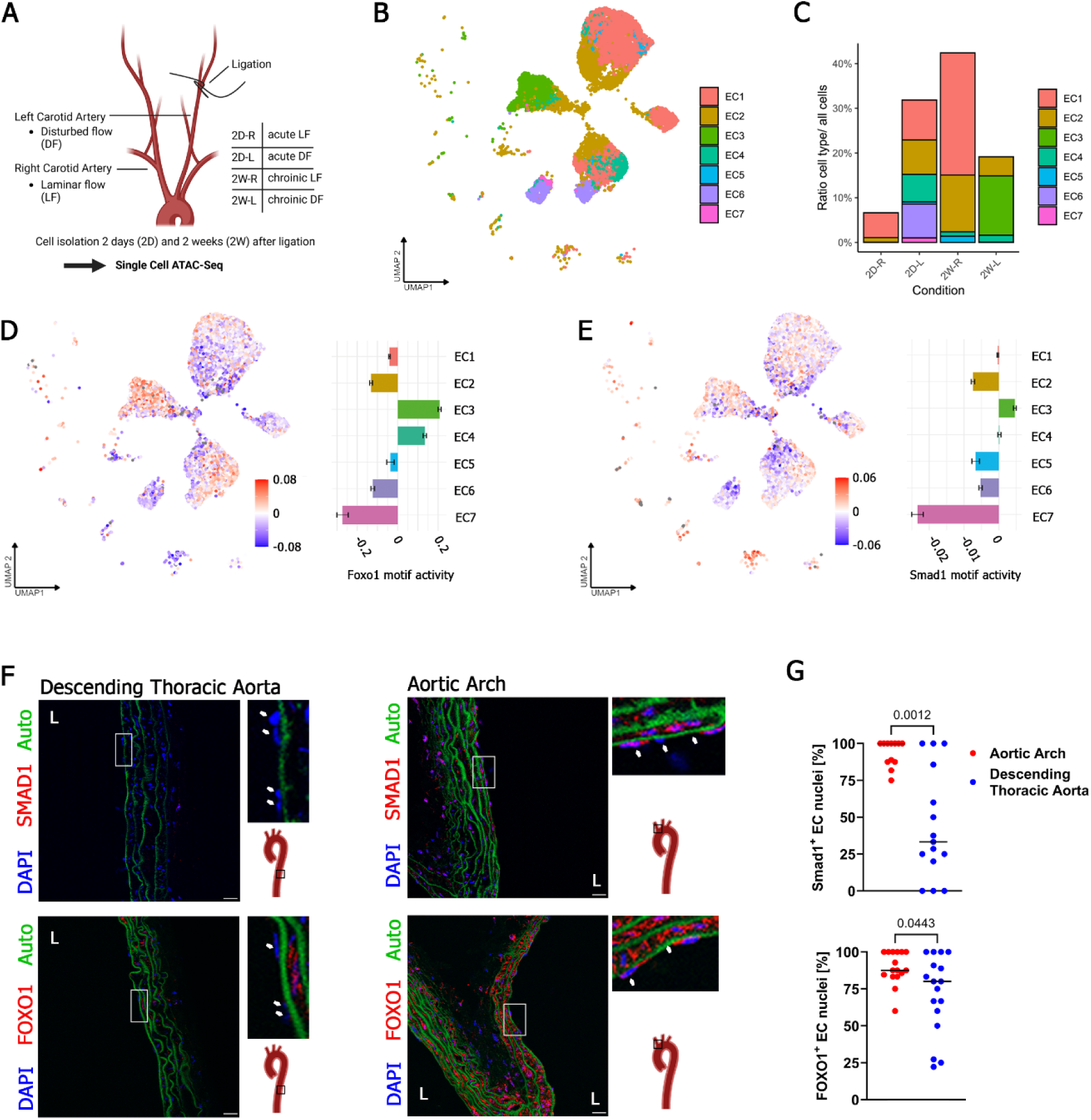
FOXO1 and SMAD1 are active in ECs exposed to disturbed flow *in vivo.* **A** Schematic depiction of carotid ligation mouse model and cell harvest strategy for scATAC-Seq samples available from PRJNA646233. **B** UMAP dimensionality reduction plot depicting identified endothelial cell (EC) clusters from scATAC-Seq data. **C** Proportions of each EC cluster compared to all cells per condition. **D** Activity of FOXO motif (JASPAR MA0480.1) amongst EC clusters on UMAP plots (upper) or bar plots (lower). **E** Activity of SMAD motif (JASPAR MA0535.1) amongst EC clusters on UMAP plots (upper) or bar plots (lower). **F** Confocal images of FOXO1 and SMAD1 immunostainings in the aortic arch or thoracic aorta of wild-type C57BL/6 mice (n=3). White arrows indicate EC nuclei. Four consecutive sections were stained and imaged per mouse. Autofluorescent (auto) signal in green originates from elastin lamellae. Scale bars are 20 µm. **G** Quantification of SMAD1 or FOXO1 positive EC nuclei of stainings from **F**. Statistics were calculated using a Mann-Whitney-U test. The Lumen is indicated by a white L.

Since FOXO1 signaling has recently been shown to be enhanced upon vascular injury and atherosclerosis^18, 19^ and FOXO1 has been demonstrated as a SMAD co-factor in a different context^21^, we next sought to determine whether FOXO1 and SMAD1 have a nuclear distribution in ECs exposed to disturbed flow in mice. For this, we performed immunofluorescence staining of either SMAD1 or FOXO1 in the descending thoracic aorta or aortic arch of mice, the latter representing chronic DF exposure **(Figure 2F**). We observed strong signals for both SMAD1 and FOXO1 in nuclei of aortic arch ECs while ECs of the descending thoracic aorta showed only weak signals (**Figure 2G**). Secondary antibody staining controls confirmed the specificity of the staining (**Figure S2**).

In summary, we showed that FOXO1 and SMAD1 proteins have a nuclear localization ECs experiencing disturbed flow, and that SMAD and FOXO motifs show the highest activity in ECs exposed to DF *in vivo*.

### SMAD1 and FOXO1 form complexes in response to atheroprone flow

Next, we investigated whether FOXO1 and SMAD may function as co-factors mediating the atheroprone flow responses of SMADs in human cells. To test this, we performed endogenous co-immunoprecipitation of either SMAD1 or SMAD2/3 with FOXO1 in HAoECs. We observed that both SMAD1 and SMAD2/3 interact with FOXO1 (**Figure 3A**). To confirm this interaction in atheroprone flow conditions we used a pneumatic pump system to apply OS (5 dyn/cm^2^, 1Hz, changing directions, atheroprone) on HAoECs for 24 h and used PS as a control (20 dyn/cm^2^, 1Hz, unidirectional, atheroprotective) (**Figure 3B**). First, to evaluate if OS induces inflammatory activation of ECs as expected for atheroprone flow, we performed a monocyte adhesion assay comparing THP-1 cell adhesion to HAoECs pre-exposed to OS or PS. We observed significantly more adhesion of monocytes to the EC monolayer in OS compared to PS (**Figure 3C,D**). Next, we showed that FOXO1 is located in the nucleus of OS-exposed HAoECs, indicating activation upon atheroprone flow exposure (**Figure S3A**). We then applied proximity ligation assay (PLA) of SMAD1 or SMAD2/3 and FOXO1 on flow-exposed HAoECs. We found that FOXO1 and SMAD1 or SMAD2/3 (**Figure 3E, Figure S3B**) showed association in both OS and PS and that this association was significantly enhanced in OS compared to PS for SMAD1-FOXO1 but not for SMAD2/3-FOXO1 (**Figure 3F, Figure S3C**). Control samples incubated with secondary probes only did not result in PLA events, highlighting the specificity of this approach (**Figure S3D**).

**Figure 3.**
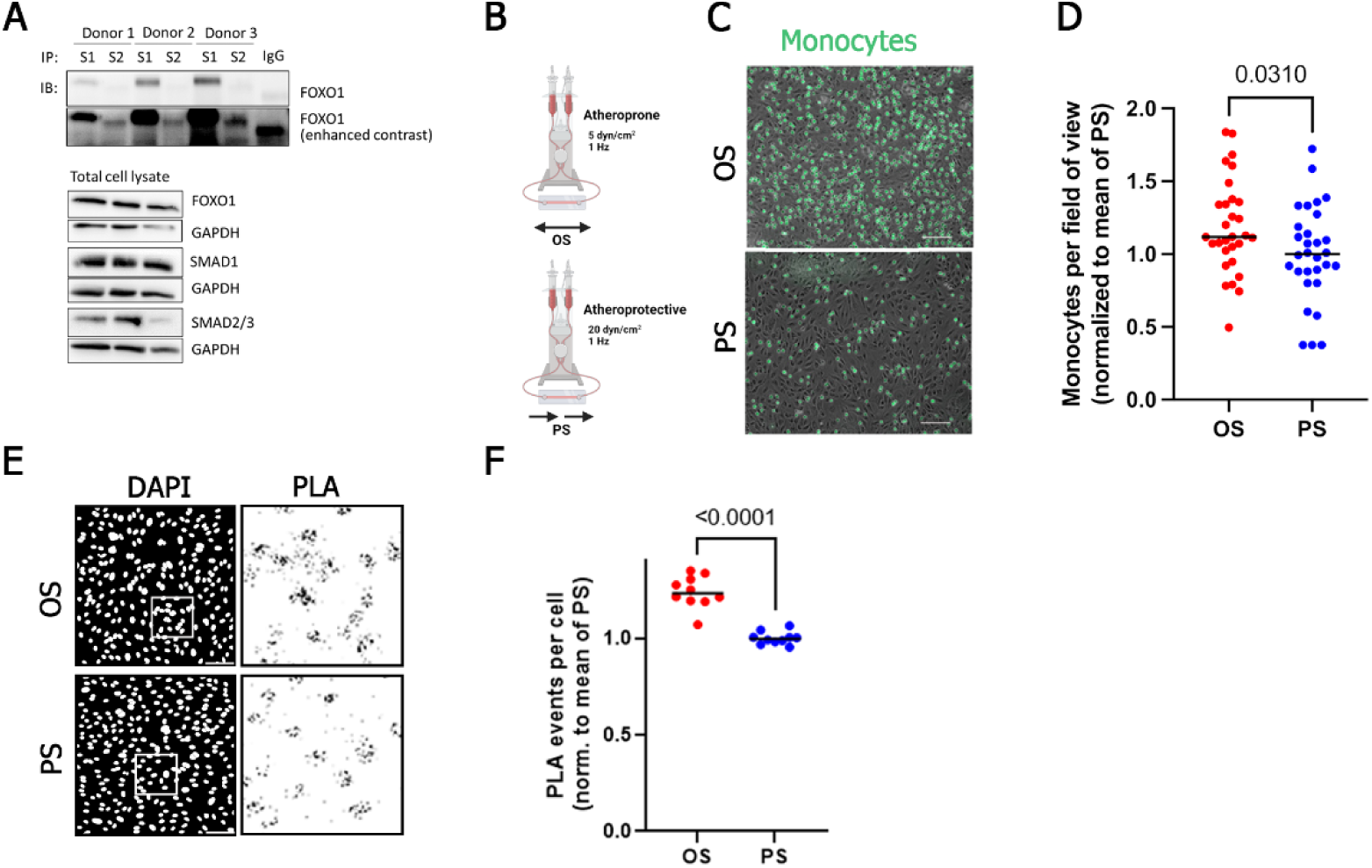
FOXO1 and SMAD1 interact upon atheroprone flow exposure of human ECs. **A** Images showing western blots from co-immunoprecipitation of total SMAD1 (S1) or total SMAD2/3 (S2) with FOXO1. Total cell lysate represents cell lysate before immunoprecipitation. IP – Immunoprecipitation, IB-Immunoblot. **B** Scheme of pneumatic pump system used to apply oscillatory (OS) and pulsatile (PS) flow on human aortic endothelial cells (HAoECs). **C** Monocyte adhesion assay using THP-1 cells labeled in green on HAoEC monolayers exposed to OS or PS (n=3). Scale bars are 100 µm. **D** Quantification of adhered monocytes from **C**. Quantification from 10 images of 3 biological donors. Statistics were calculated using a two-tailed Student’s *t*-test. **E** Confocal images of SMAD1-FOXO1 interaction assessed by proximity ligation assay (PLA) on HAoECs exposed to OS or PS (n=2). Scale bars are 100 µm. **F** Quantification of PLA events per cell from **E**. Quantification from 10 images of 2 biological donors. Statistics were calculated using a two-tailed Student’s *t*-test.

Together, we identified FOXO1 as an atheroprone flow-dependent SMAD1 co-factor.

### SMAD and FOX motifs are enriched in regions associated with enhanced transcription

To investigate whether FOXO1 and SMADs indeed co-regulate transcription upon atheroprone flow exposure in humans, we performed unbiased ATAC-Seq and RNA-Seq of HAoECs exposed to OS and PS which allowed us to correlate DARs with increased or decreased expression of assigned genes. Using the RNA-Seq data, we found that OS and PS led to the upregulation of 305 and 264 genes, respectively (adjusted p-value < 0.05, |log2(fold change)| > 0.585). Here, OS led to the upregulation of SMAD target genes *EDN1*, *CCN2*, and *ID3* while PS upregulated known atheroprotective genes *KLF2*, *KLF4*, and *TEK* (**Figure 4A**). GO enrichment of KEGG pathway terms identified an overrepresentation of genes belonging to the PI3K pathway, which is upstream and inhibitory to FOXO1 action, and to be down-regulated in OS (**Figure 4B**) ^67, 68^. By analyzing the ATAC-Seq data, we identified 949 and 2,942 more accessible DARs in OS and PS, respectively (adjusted p-value < 0.05, |log2(fold change)| > 0.585) (**Figure 4C**). Amongst regions that gained accessibility upon OS were those associated with SMAD target genes *EDN1*, *CCN2*, *SERPINE1*, and *ID3*. In contrast, regions associated with atheroprotective genes *KLF2*, *KLF4*, *TEK*, and *CYP1B1* showed enhanced accessibility in PS, in line with our RNA-Seq data (**Figure 4C**). To validate enhanced SMAD and FOXO activity upon atheroprone OS, we next performed TF footprinting analysis using TOBIAS^47^. We found footprints of both SMAD binding and FOXO motifs (**Figure 4D**) to be enriched in OS samples. To further investigate if chromatin accessibility changes are causative for changes in gene expression in atheroprone flow, we integrated RNA-Seq and ATAC-Seq data, *i*.*e*. identified genes that show concomitant accessibility and expression changes. We found that 32 DARs were associated with enhanced expression of the nearest associated gene in OS compared to 96 in PS (**Figure 4E**). Amongst those genes upregulated with an accompanying opening region in OS was SMAD target gene *EDN1*. To validate that SMADs and FOXO1 drive concomitant accessibility and expression changes in OS we performed motif enrichment analysis of the regions that were positively correlated with enhanced gene expression and indeed detected SMAD and FOX motifs, suggesting shared transcriptional regulation by FOXO and SMADs in human cells (**Figure 4F**).

**Figure 4.**
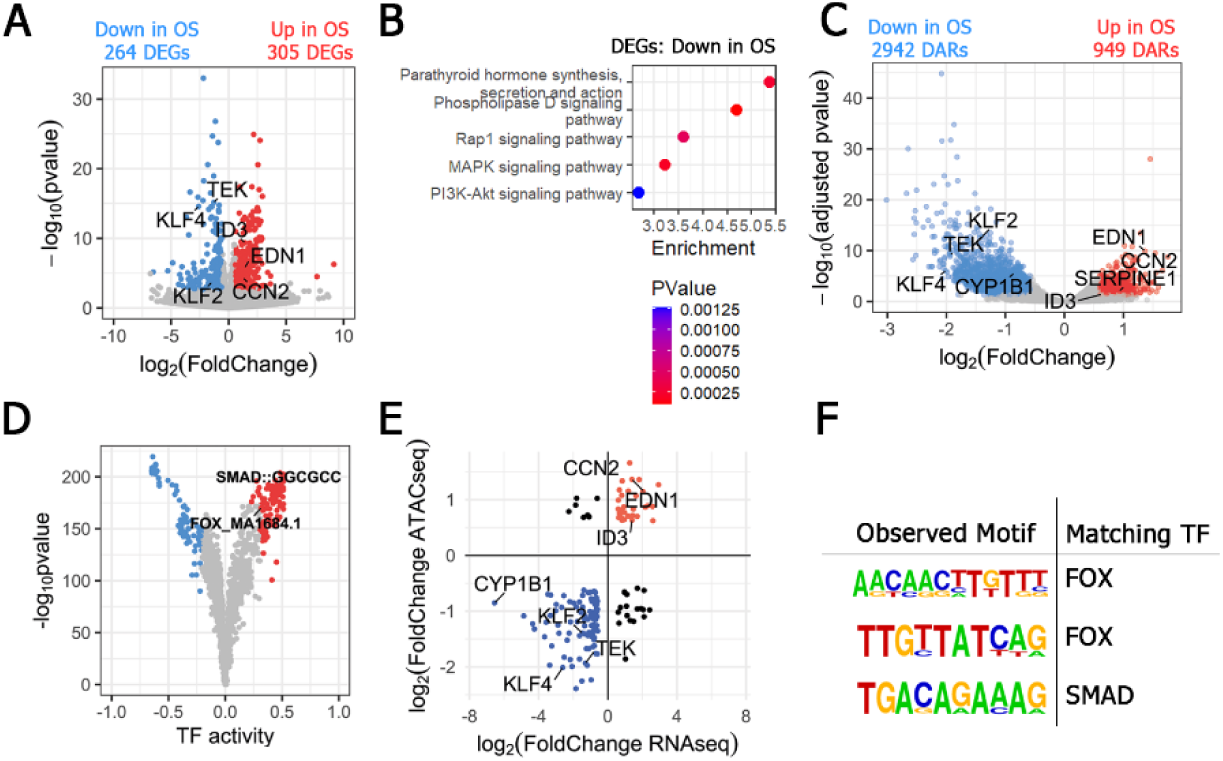
Shared FOXO and SMAD motif enrichment in atheroprone flow exposed human aortic endothelial cells (HAoECs). **A** Volcano plot of differentially expressed genes (DEGs) (adjusted p-value < 0.05, fold change > |log_2_(0.585)|) from RNA-Seq data of oscillatory (OS) and pulsatile (PS) flow exposed HAoECs (n=3). **B** Gene ontology enrichment analysis of genes down regulated by OS. **C** Volcano Plot of differentially accessible regions (DARs) (adjusted p-value < 0.05, fold change > |log_2_(0.585)|) from ATAC-Seq data of OS and PS exposed HAoECs (n=3). **D** Footprinting analysis using DARs from ATAC-Seq data. Transcription factors (TFs) in color have -log10(p-value) above the 95% quantile and/ or differential binding scores smaller/larger than the 5% and 95% quantiles (top 5% in each direction). Motifs for SMAD or FOX TF members are highlighted. **E** Integration of DEGs and DARs from RNA-Seq/ ATAC-Seq. **F** Motif enrichment analysis of DARs that show enhanced expression of the associated gene. Found motifs were screened for similarities to motifs of SMADs or known SMAD co-factors.

### SMADs and FOXO1 co-regulate atheroprone gene expression

To further investigate the mechanistic background of SMAD-FOXO transcriptional co-regulation, we analyzed differentially accessible loci that display both Smad binding motifs (SBEs) and FOX binding sites. We found that out of all regions that show significantly increased accessibility in OS, 51.2 % were positive for SMAD and FOXO binding sites (**Figure 5A**) highlighting FOXO TFs as possible co-regulators of SMAD-induced gene transcription. Next, we asked which of the SMAD/FOXO double-positive regions correlate with enhanced expression of assigned genes. We found 28 regions, 3 of which were associated with SMAD target genes *EDN1* (2x) and *ID3* (**Figure 5B**). By visualizing the promoter region of *EDN1*, we found that FOXO and SMAD binding sites are located in close proximity (**Figure 5C**), suggesting concerted action of SMAD1 and FOXO TFs. To confirm this, we tested the responsivity of the FOXO binding site with one adjacent SMAD binding site and the corresponding linker sequence from the *EDN1* promoter to FOXO1 and SMADs using a luciferase-based reporter gene assay in HEK293T cells (**Figure 5D**). We could show that the tested *EDN1* promoter sequence is sensitive to FOXO1, and this sensitivity is lost upon mutation of the FOXO binding site (**Figure 5D**). However, we did not observe sensitivity to SMAD activation upon BMP6 stimulation (**Figure S4A**) suggesting a more complex SMAD-dependent regulation of *EDN1* involving several SMAD binding sites. To test whether SMAD and FOXO are nonetheless co-regulating gene expression, we constructed a synthetic reporter gene plasmid containing two SMAD binding sites and a FOXO binding site (**Figure 5E**). Using FOXO1 overexpression^30^ and BMP6 stimulation we could show that this sequence is sufficient to induce transcription based on BMP stimulation (∼ 4-fold) or FOXO1 overexpression (∼3.8-fold). Furthermore, the induced gene expression was even higher when both, FOXO1 overexpression and BMP6 stimulation, were performed simultaneously (∼ 5.8-fold), highlighting a synergistic regulation of transcription by SMADs and FOXO1 (**Figure 5E**). To additionally interrogate shared binding of SMAD and FOXO1, we used Cleavage Under Targets & Release Using Nuclease followed by sequencing (CUT&RUN-Seq) to evaluate binding of SMAD1 and SMAD2/3 in HAoECs and public ChIP-seq data from human umbilical vein ECs (HUVECs) for FOXO1 to confirm shared SMAD and FOXO1 binding in the promoter regions of *ID3* and *EDN1* (**Figure 5F**). To further investigate if FOXO1 indeed co-regulates transcription of SMAD target genes in response to atheroprone flow we performed siRNA-mediated KD of FOXO1 in HAoECs. First, we confirmed the efficient reduction in *FOXO1* mRNA levels (**Figure S4B**). We then subjected HAoECs to OS and observed that KD of FOXO1 led to decreased expression of SMAD target genes *ID3* and *EDN1* (**Figure 5G**).

**Figure 5.**
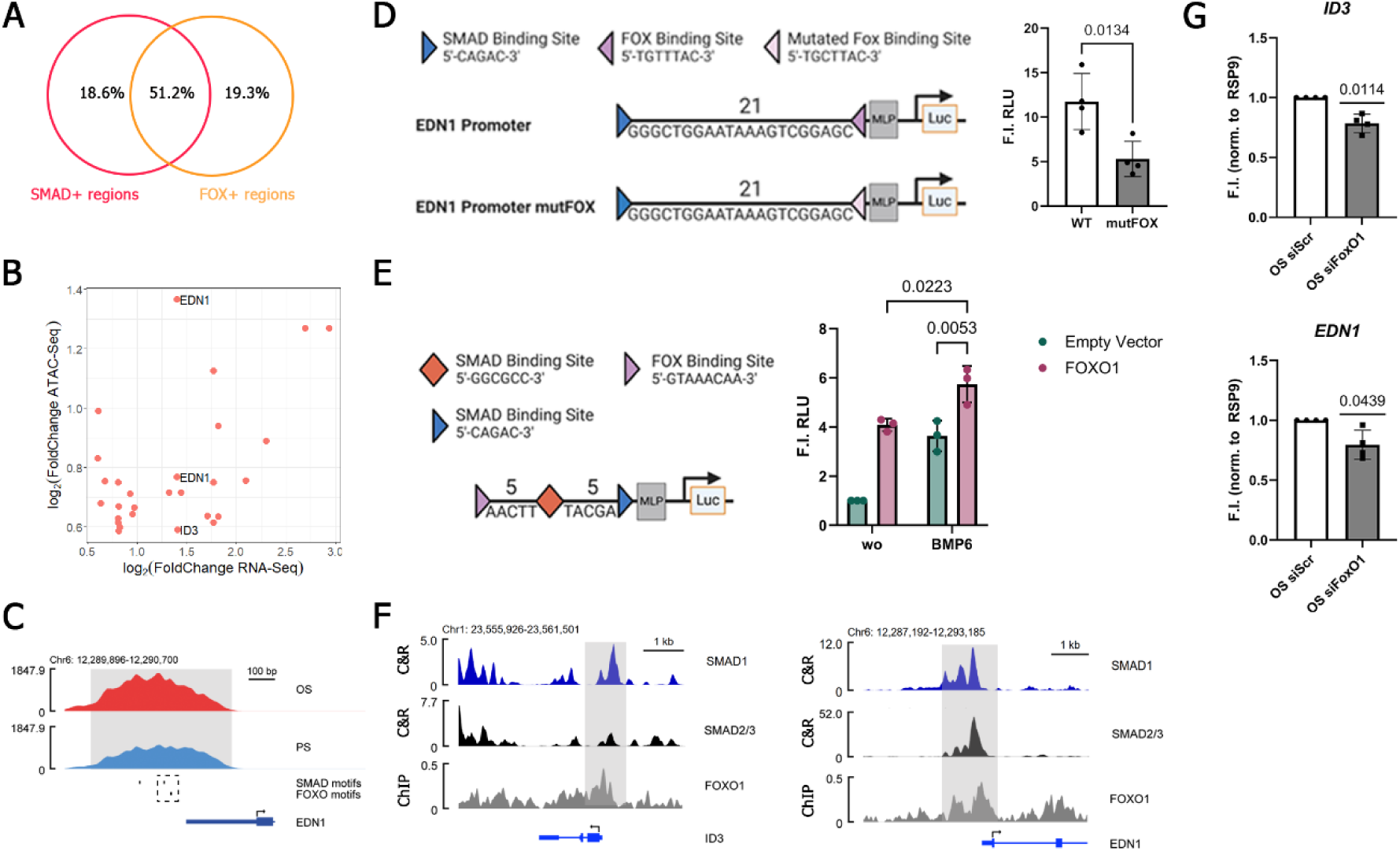
Shared transcriptional regulation by FOXO1 and SMADs. **A** Venn diagram showing the percentage of differentially accessible ATAC-Seq regions (DARs) (adjusted p-value < 0.05, fold change > |log_2_(0.585)|) positive for SMAD and/ or FOXO motifs in human aortic endothelial cells (HAoECs) (n=3). **B** SMAD and FOXO-positive regions which are associated with genes that display enhanced expression. BMP target genes are highlighted. **C** Genome browser view of the *EDN1* promoter displaying ATAC-Seq data from HAoECs exposed to oscillatory (OS) or pulsatile (PS) flow. SMAD and FOXO motifs are highlighted below. The dashed line indicates regions used in **D**. Gray shading highlights areas of interest. **D** Reporter gene assay in HEK293T cells using a genomic region of the *EDN1* promoter region indicated by a dashed line in **C**. Values are fold inductions of FOXO1 overexpression compared to an empty vector control (n=4). Quantification shows the mean of experiments +/- standard deviation. Statistics were calculated using a two-tailed Student’s *t*-test. **E** Reporter gene assay of a synthetic SMAD and FOXO motif containing plasmid in HEK293T cells (n=3). Values are normalized to the unstimulated empty vector control and represent means of 3 independent experiments +/- standard deviation. Statistics were calculated using a two-way ANOVA with Šidák’s post-hoc test. **F** Genome Browser view of SMAD1 and SMAD2/3 CUT&RUN (C&R) of BMP9 stimulated HAoECs (n=2 each) and FOXO1 ChIP-Seq in human umbilical vein ECs upon overexpression of constitutively active FOXO1 (GSE128635). Gray shading highlights area of interest. **G** qPCR analysis of BMP target genes in HAoECs exposed to OS upon siRNA-mediated depletion of FOXO1. Values are normalized to the scrambled control and are depicted as mean +/- standard deviation from 4 independent experiments. Statistics were calculated using one sample *t*-test for siFOXO1.

In conclusion, we found that FOXO1 and SMADs synergistically co-regulate transcription in human cells, bind to the same genomic loci in response to atheroprone flow and bind promoter regions of shared target genes.

### FOXO1 and SMADs co-regulate EndoMT via *SNAI2*

SMADs are main inducers of EndoMT^69^ and it has recently been shown that SMADs induce EndoMT downstream of DF^70^. Since FOXO1 is co-regulating SMAD dependent gene expression in response to atheroprone flow, we then asked if FOXO1 is involved in EndoMT regulation. We thus performed RNA-Seq of HAoECs subjected to OS in the presence of AS1842856 (0.5 µM), which blocks FOXO1 DNA binding (**Figure 6A**)^71^. We investigated the expression of EndoMT-associated genes and observed that their expression decreased significantly upon FOXO1 inhibition (**Figure 6B**). Interestingly, when performing GO enrichment analysis, terms corresponding to SMAD and PDGF pathways, both known regulators of EndoMT, were found (**Figure 6C**). Moreover, KEGG pathway analysis revealed inflammation (TNF, NF-κB, p53) and atherosclerosis-related pathways in addition to the FOXO signaling pathway (**Figure S6A**). We then asked whether FOXO1 mediates the expression of EndoMT genes by opening chromatin which can be subsequently bound by other TFs like SMADs. We thus performed ATAC-Seq in the presence of AS1842856 (0.5 µM) (**Figure 6A**). Intriguingly, we observed changes in chromatin accessibility only at promoters of *PDGFA* and *PDGFB* (**Figure 6D**), while those of the other regulated EndoMT-related genes did not change (**Figure 6E**). We thus speculated that FOXO1 upregulates the expression of EndoMT TFs which subsequently induces expression of other EndoMT target genes. We observed that accessibility of the promoter region of *SNAI2* (encoding SLUG) showed only minor changes upon FOXO1 inhibition (**Figure 6F**). However, we found that overexpression of constitutively active FOXO1 (FOXO1-CA) in HUVECs leads to an increase in histone 3 lysine 27 acetylation (H3K27Ac), a mark for active chromatin (**Figure 6F**). Moreover, we used existing FOXO1 ChIP-Seq data in HUVECs and CUT&RUN-Seq of SMAD1 and SMAD2/3 in HAoECs and could show that both SMADs and FOXO1 bind the *SNAI2* promoter region (**Figure 6F**).

**Figure 6.**
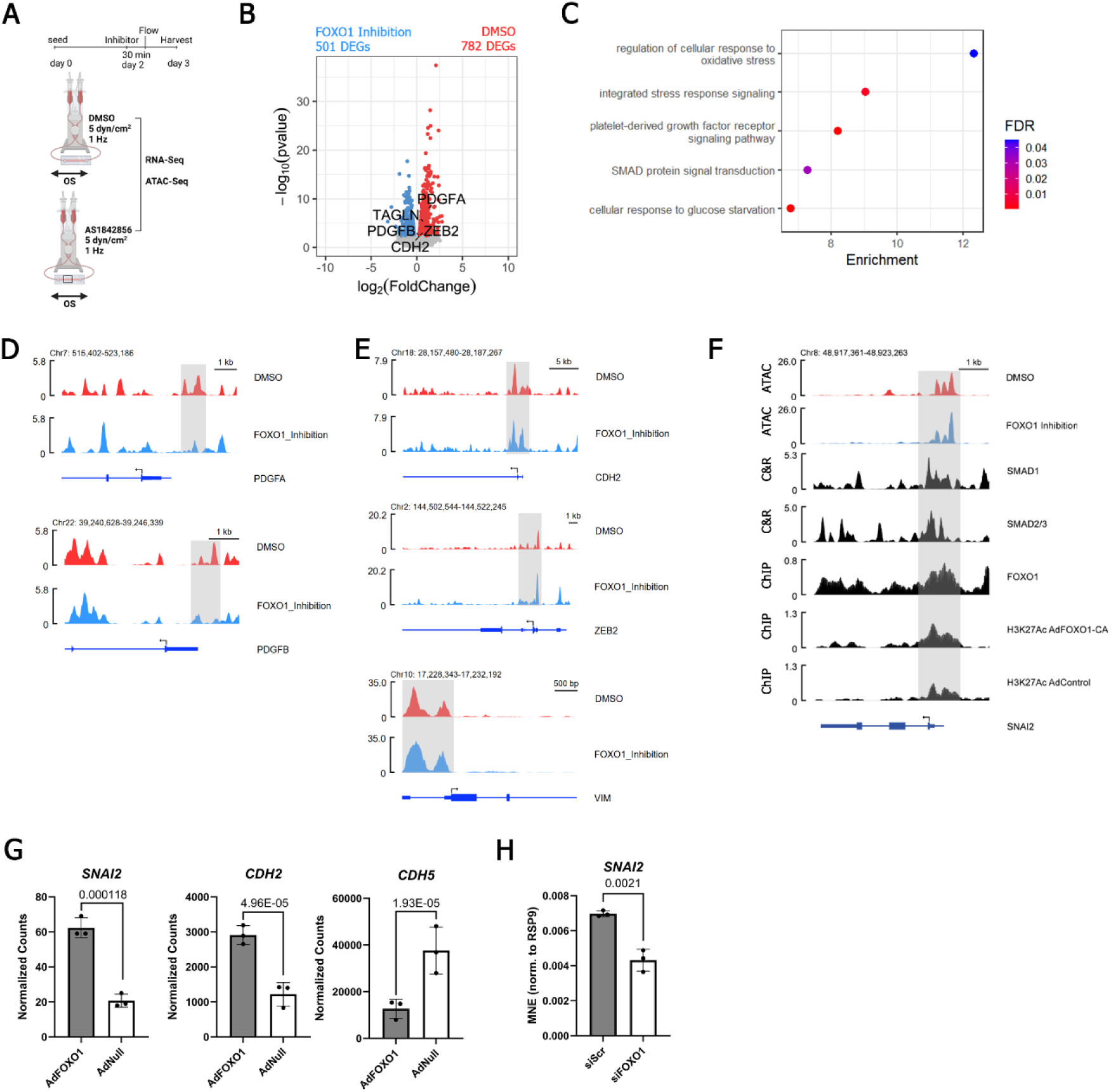
FOXO1 mediates endothelial-to-mesenchymal transition (EndoMT) in atheroprone flow-exposed human aortic endothelial cells (HAoECs). **A** Schematic overview of the experimental setup. **B** Volcano plot of differentially expressed genes (DEGs) (adjusted p-value < 0.05, fold change > |log_2_(0.585)|) from RNA-Seq data of HAoECs exposed to oscillatory flow (OS) in absence or presence of FOXO1 inhibitor AS1842856 (0.5 µM) (n=3). Genes corresponding to EndoMT are highlighted. **C** Gene ontology enrichment analysis of DEGs corresponding to active FOXO1. **D**,**E** Genome browser views of ATAC-Seq data from HAoECs exposed to OS in the presence or absence of FOXO1 inhibitor AS1842856 (0.5 µM) (n=3). Gray shading highlights areas of interest. **F** Genome browser view of ATAC-Seq data from HAoECs exposed to OS or pulsatile flow (PS), SMAD1; SMAD2/3 CUT&RUN of BMP9 stimulated HAoECs, and ChIP-Seq of FOXO1 and histone 3 lysine 27 acetylation (H3K27Ac) in human umbilical vein ECs (HUVECs) upon overexpression of constitutively active FOXO1 (FOXO1, H3K27Ac) or an empty control vector (H3K27Ac) (GSE128635). Gray shading highlights areas of interest. **G** Normalized counts from RNA-Seq of HUVECs transduced with control vector or constitutively active FOXO1 (GSE128636). Displayed p-values were calculated using DESeq2 (adjusted p-value) to account for multiple testing and non-normality of RNA-Seq data. Wald test p-values are corrected for multiple testing using the Benjamini and Hochberg method. **H** RT-qPCR analysis of HAoECs transfected with control or FOXO1 siRNA (n=3). Values are depicted as mean normalized expression +/- standard deviation. Statistics were calculated using a two-tailed Student’s *t*-test.

To further corroborate that FOXO1 regulates *SNAI2* transcription in ECs we analyzed existing RNA-Sequencing data of FOXO1-CA overexpression in HUVECs^72^. We observed a strong upregulation of *SNAI2* expression in FOXO1-CA samples compared to the control vector (**Figure 6G**). Moreover, the expression of *CDH5* (VE-Cadherin) decreased while expression of *CDH2* (N-Cadherin) increased, indicating a partial loss of endothelial identity and gain of a mesenchymal signature upon FOXO1-CA overexpression (**Figure 6G**). We could furthermore show that silencing of *FOXO1* in HAoECs leads to decreased expression of *SNAI2* (**Figure 6H**).

Overall, we showed that FOXO1 and SMAD1 regulate EndoMT in response to atheroprone flow through upregulation of *SNAI2*.

### FOXO1 and SMAD1 are found in atherosclerotic lesions in mice

Finally, we investigated the presence of FOXO1 and SMAD1 in atherosclerotic lesions using aged LDLR knock-out mice, which exhibited established atherosclerotic plaques (**Figure 7A**). Immunofluorescence staining of SMAD1 and FOXO1 was performed on sections of the aortic arch containing these plaques. Our analysis revealed that both FOXO1 and SMAD1 were present in the nuclei of ECs, supporting a collaborative role for FOXO1 and SMAD1 in atherosclerotic lesions.

**Figure 7.**
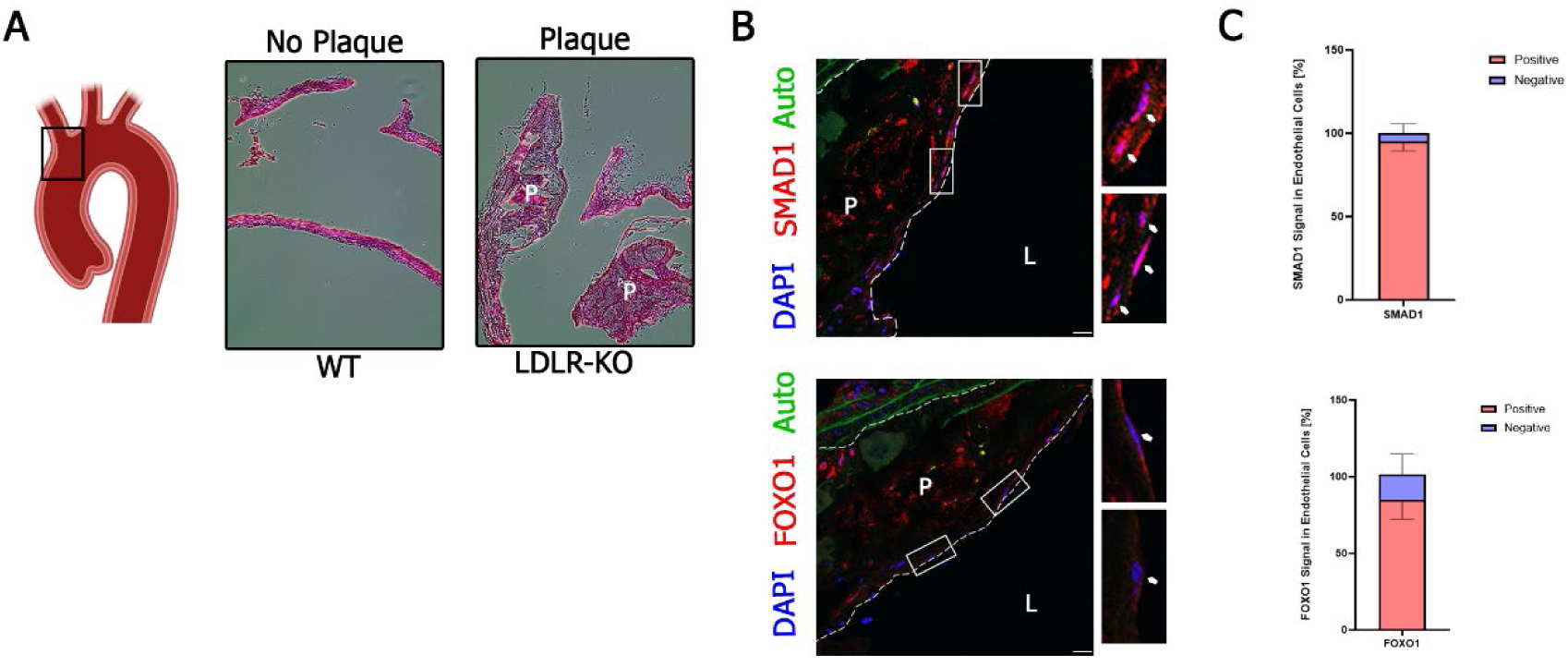
FOXO1 and SMAD1 are active in endothelial cells (ECs) in atherosclerotic lesions. **A** Hematoxylin and Eosin staining of wild-type (WT) aortic arch or of atherosclerotic plaques in the aortic arch of a low-density lipoprotein receptor (LDLR) knock-out mouse (n=3, 3-5 sections analyzed per mouse). **B** Confocal images of FOXO1 and SMAD1 immunostainings on atherosclerotic lesions of aged LDLR knock-out mice (n=3). Four consecutive sections were stained and imaged per mouse. White arrows indicate EC nuclei. Autofluorescent (auto) signal in green originates from elastin lamellae. Scale bars are 20 µm. **C** Quantification of SMAD1 or FOXO1 positive EC nuclei compared to the total number of EC nuclei of staining from **B**. Values are depicted as mean +/- standard deviation. Lumen is indicated by a white L. Plaque is indicated by a white P.

## Discussion

We have previously shown that BMP signaling is highly dependent on the mechanical context of cells, on the level of receptors and co-receptors^7^, SMAD phosphorylation and co-factor binding^73, 74^, or regulation of chromatin accessibility and gene expression^9^. In the current study, we extended this knowledge by identifying a novel role for FOXO1 in ECs, where it facilitates EndoMT in conjunction with SMADs (**Figure 8**). Mechanistically, upon exposure to atheroprone flow, FOXO1 and SMADs form complexes that bind to the *SNAI2* promoter, leading to increased H3K27 acetylation and enhanced gene expression (**Figure 8**).

**Figure 8.**
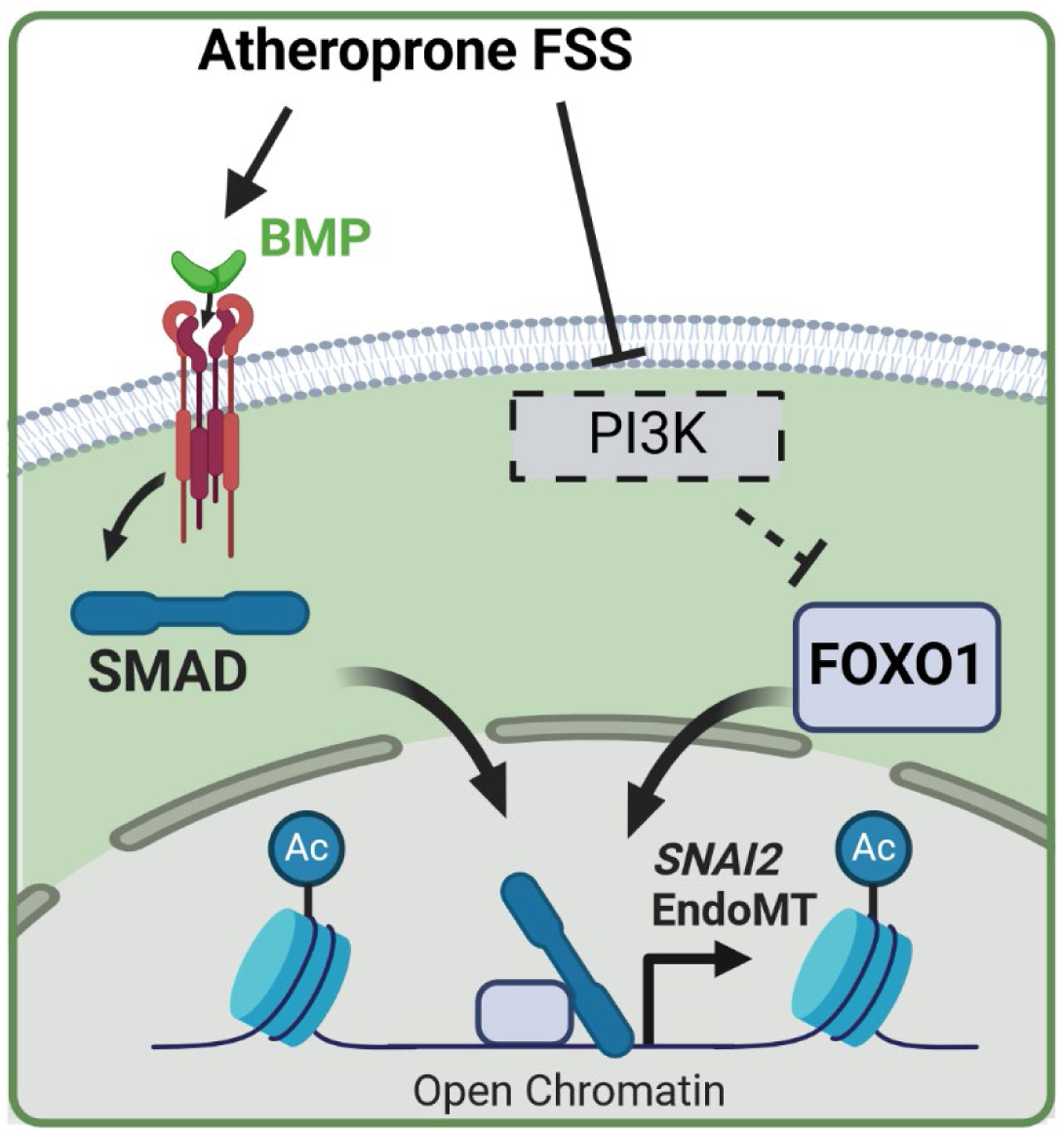
Graphical summary-FOXO1 and SMADs regulate EndoMT in response to atheroprone flow by inducing the expression of *SNAI2*.

Generally, little is known about the contribution of FOX TFs to EndoMT. In a recent study, authors investigated EC and mesenchymal marker gene expression upon TGFβ stimulation and knock-down or pharmacological inhibition of FOXM1 to state that FOXM1 is a critical driver of EndoMT downstream of TGFβ in HUVECs^75^. EndoMT and epithelial-to-mesenchymal transition (EMT) are distinct, yet closely related processes, sharing similar steps and TFs involved^76^. Interestingly, several FOX-TFs have demonstrated roles in promoting EMT in cancer, including FOXM1, FOXS1, FOXQ1, and FOXC1^77-83^. Conversely, FOXO1 has been shown to inhibit EMT and cancer cell invasion^84^. This depicts an interesting contrast to our data and highlights the differences between EMT and EndoMT and the importance of the cellular system used. In general, the existing data highlights the involvement of several FOX family TFs in EMT and suggests that members of this TF family have unexplored roles in regulating EndoMT.

While previous reports have documented interactions between FOX proteins and SMAD3 or SMAD4^21, 85, 86^, our study expands this knowledge by demonstrating interactions between FOXO1 and both SMAD1 and SMAD2/3. These dual interactions may result from FOXO1’s ability to bind to the common SMAD, SMAD4, which subsequently forms complexes with active SMADs. Notably, we observed differences in the complex formation of FOXO1-SMAD1 compared to FOXO1-SMAD2/3 in response to atheroprone flow, despite both SMAD1 and SMAD2/3 being activated by such flow conditions. This suggests the involvement of distinct, yet unidentified co-factors, that may facilitate or stabilize FOXO1-SMAD1 complex formation.

In our previous study, we hypothesized that SMAD-dependent transcriptional regulation is facilitated by pioneer factors that open chromatin regions typically inaccessible for SMADs and other TFs^9^. While FOXO1 exhibits pioneering capabilities^22^, our findings indicate that its DNA binding is not essential for chromatin opening in the promoter regions of EndoMT genes in response to atheroprone flow. Recent studies on cellular reprogramming highlight the existence of chromatin regions enriched in somatic TF motifs, which can sequester these factors in a sponge-like manner^87^. These regions become accessible through the action of pioneering or reprogramming TFs, enabling cell fate transitions^87^. Therefore, we speculate that FOXO1 may contribute to the opening of these sponge-like regions in response to atheroprone flow, leading to the sequestering of endothelial lineage TFs and subsequently promoting EndoMT.

We found a down-regulation of genes corresponding to the GO term “PI3K-Akt signaling pathway” in OS compared to PS control, in line with reports that showed PI3K/ AKT signaling to be enhanced in laminar FSS-exposed ECs^67^. Furthermore, PI3K/ AKT signaling is upstream of FOXO phosphorylation and thus inactivation and cytoplasmic retention^20^. Hence, it is likely that in addition to lower expression of FOXO1 in PS, FOXO1 is also deactivated by enhanced PI3K/ AKT signaling, which leads to attenuation of FOXO1-mediated gene regulation^20^.

Additionally, experiments in mice that bear constitutively deacetylated alleles of FOXO1 (*Foxo1^KR/KR^*) indicate the involvement of FOXO1 in the development of atherosclerosis^15^. In ECs of *FOXO1^KR/KR^* mice, FOXO1 preferentially locates to the nucleus and induces expression of inflammatory genes^15^. In addition, *Foxo1^KR/KR^* mice bred on an *LDLR^-/-^* background show increased atherosclerotic lesion formation compared to *LDLR^-/-^*;*FOXO1^WT^*controls ^15^. Moreover, FOXO1 was found to mediate a pro-inflammatory response in response to a metabolite stimulus involved in atherosclerosis^18^. Furthermore, mice lacking all FOXO isoforms (FOXO1, FOXO3A, FOXO4) show less atherosclerotic lesion formation in the *LDLR^-/-^* background^16^. In humans, enhanced FOXO1 expression was found in calcified femoral arteries and inhibition of FOXO1 signaling by a small molecule inhibitor prevented calcification in patient-derived fibroblasts^17^. However, knock-down of FOXO1/3 in vascular smooth muscle cells (VSMCs) led to enhanced *Runx2* expression and VSMC calcification^88^.

## Conclusions

In conclusion, our study provides new insights into the transcriptional regulation activated by atheroprone flow. Through the integration of ATAC-Seq and RNA-Seq data from primary human aortic ECs and SMAD knock-out mouse models, we identified FOXO1 as a novel co-regulator of SMAD transcriptional response induced by atheroprone flow. This regulation encompasses key atheroprone genes, such as *EDN1,* and notably influences EndoMT by upregulating *SNAI2* (encoding for SLUG). These findings highlight the potential role of FOXO1-SMAD interactions in atherosclerosis. Therefore, targeting these interactions may offer a promising strategy to specifically inhibit the atheroprone SMAD-mediated responses in ECs, ultimately addressing endothelial dysfunction and atherosclerosis.

## Novelty and Significance

### What Is Known?

- Atheroprone flow contributes to endothelial dysfunction, which is a key factor in the development of atherosclerosis, partly driven by BMP signaling
- SMAD signaling induces EC homeostasis and dysfunction and has a prominent role in EndoMT
- SMAD signaling is dependent on co-factors

### What New Information Does This Article Contribute?

- We demonstrate that FOXO1 and SMAD1 form complexes in endothelial cells, particularly in response to atheroprone flow
- Our findings reveal that FOXO and SMAD motifs are enriched in DNA loci linked to increased transcription and chromatin accessibility under atheroprone flow conditions
- Depletion of FOXO1 results in diminished expression of genes induced by atheroprone flow, including BMP target genes

The intricate mechanisms underlying atherosclerosis at focal sites in arteries exposed to atheroprone flow remain incompletely understood. Our study identifies complexes of SMAD1 and FOXO1 that are induced in these conditions and play a role in driving the pathological transcriptional response in endothelial cells both in mice and in human cells. Further research validating the importance of these complexes *in vivo* and identifying the exact sites of interaction of FOXO1 and SMAD1 will highlight the opportunities to design intervention strategies to treat or prevent atherosclerosis.

## Acknowledgements

This work was made possible by the collaboration within the Fonds voor Wetenschappelijk Onderzoek (FWO)-WOG network (WOG001420N), which supported AZ, AL, FI, PM, JJ and PK. We would like to acknowledge the assistance of the Core Facility BioSupraMol supported by the DFG.

## Supplementary Figures

**Figure S1.**
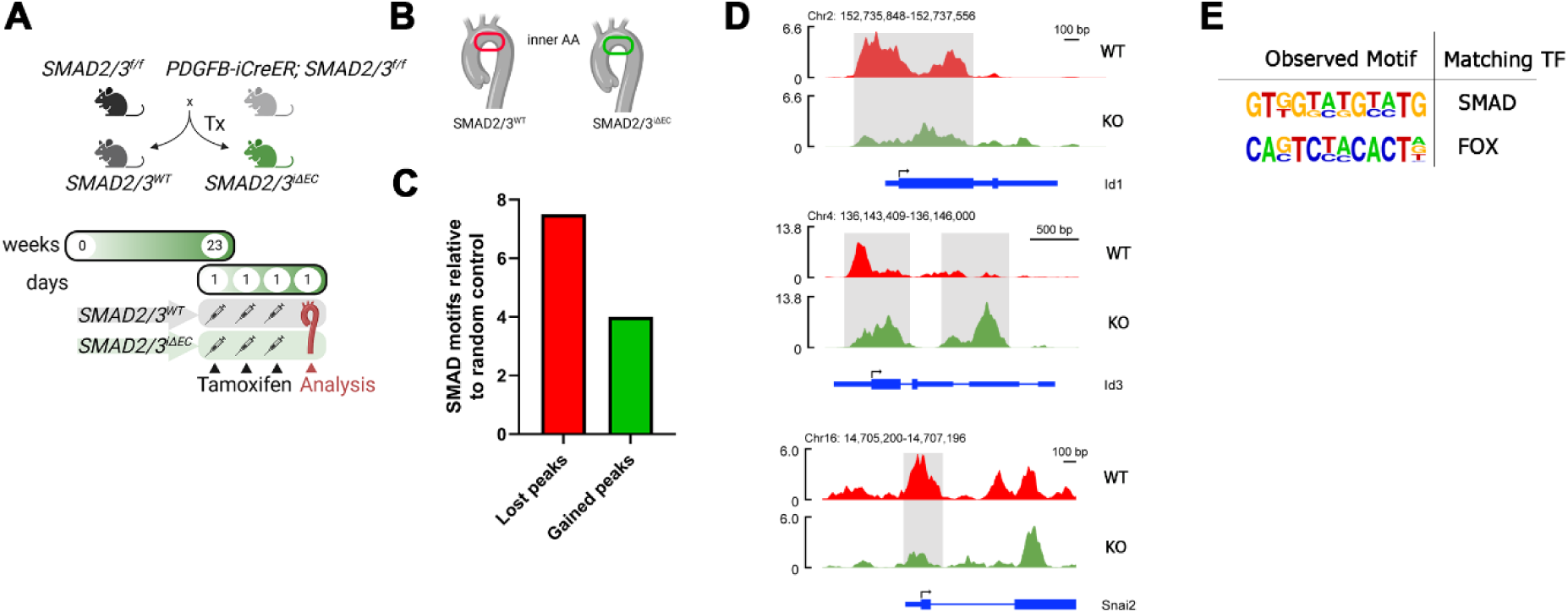
FOX motifs are enriched in SMAD2/3 dependent peaks in mouse endothelial cells (ECs). **A** Scheme of mouse breeding and tamoxifen injection. **B** Isolation of ECs from the inner curvature of the aortic arch (AA) of *SMAD2/3^WT^* (red) or *SMAD2/3^iΔEC^* (green) mice for ATAC-Seq (n=2/3 *SMAD2/3^WT^* and *SMAD2/3^iΔEC^* mice). **C** Number of SMAD motifs (*GGCGCC*) found in differentially accessible peaks (FDR<0.2) compared to random genomic regions of similar size and number. **D** Genome browser view showing ATAC-Seq tracks of ECs isolated from the inner curvature of the AA of WT (red) or SMAD2/3 knock-out (green) mice. Gray shading highlights areas of interest. **E** Motifs enriched in ATAC-Seq peaks that are lost upon SMAD2/3 knock-out identified by HOMER. Found motifs were screened for similarities to motifs of SMADs or known SMAD co-factors.

**Figure S2.**
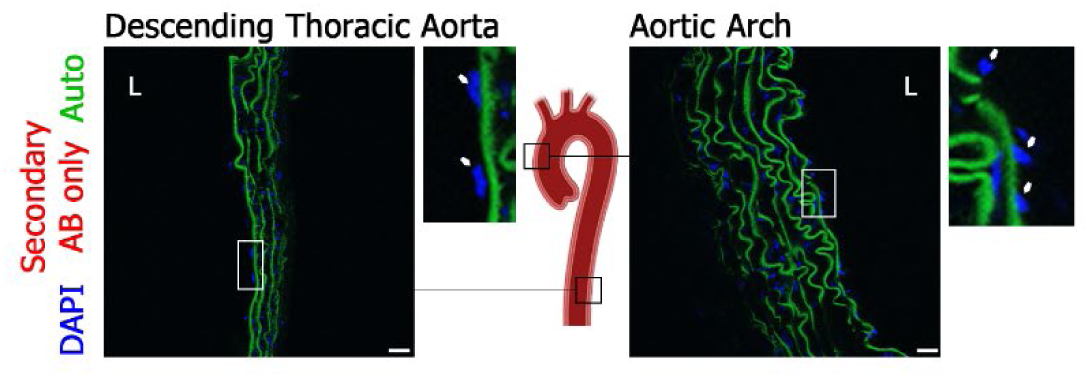
Secondary antibody control for immunostainings of mouse aortae. Confocal images of immunostainings in the aortic arch or descending thoracic aorta of wild-type C57BL/6 mice using secondary antibody only (n=3 mice). White arrows indicate EC nuclei. Autofluorescent (auto) signal in green originates from elastin lamellae. Scale bars are 20 µm.

**Figure S3.**
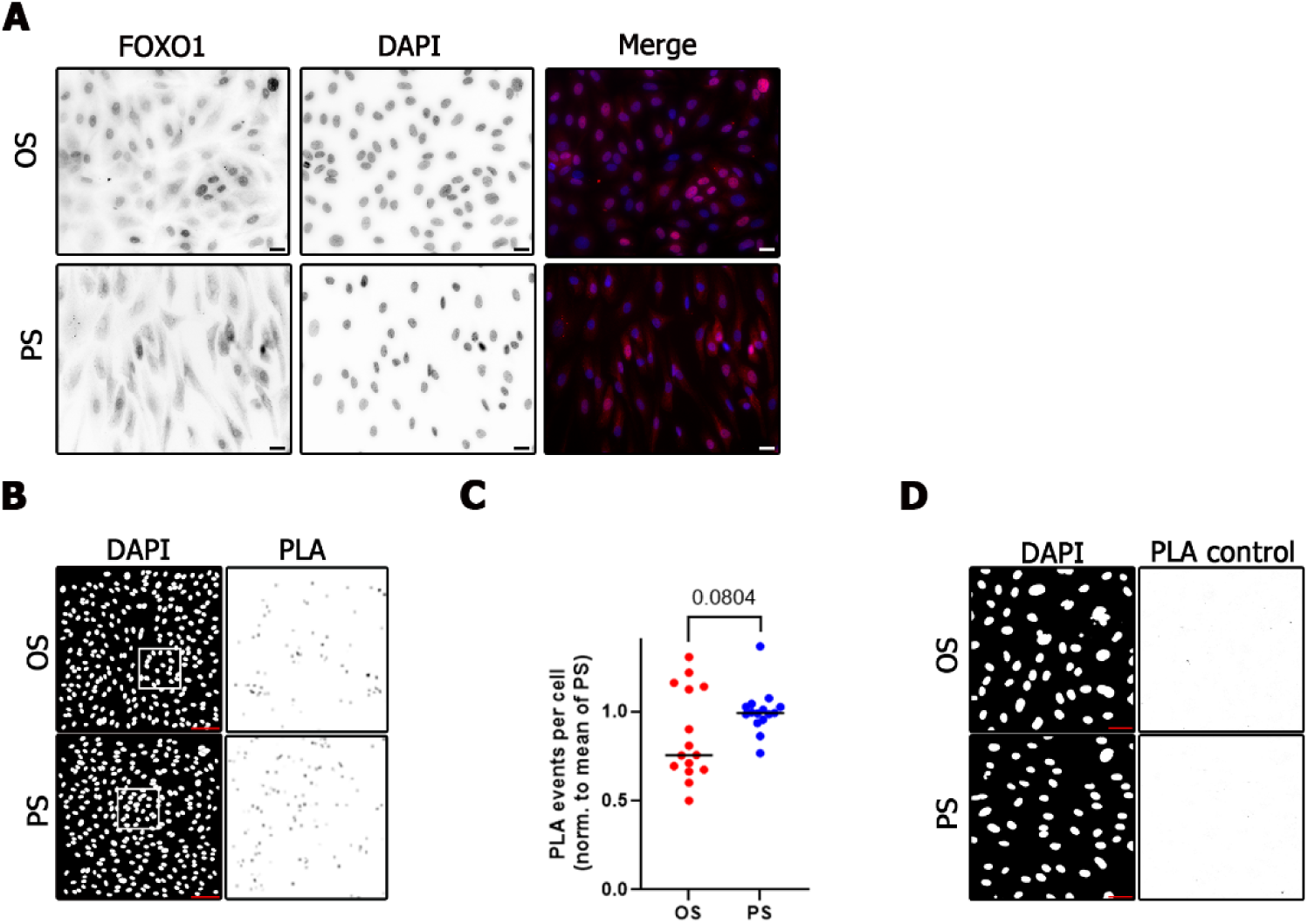
FOXO1 is active in atheroprone flow-exposed endothelial cells (ECs) and interacts with SMAD2/3. **A** Epifluorescence microscopy images of FOXO1 immunostainings (red) in human aortic ECs (HAoECs) exposed to oscillatory (OS) or pulsatile (PS) flow. Nuclei are counterstained with DAPI (blue). Representative images of 3 biological replicates with 4 images each. Scale bars are 20 µm. **B** Confocal images of SMAD2/3-FOXO1 interaction assessed by proximity ligation assay (PLA) on ECs exposed to OS or PS. Scale bars are 100 µm. **C** Quantification of PLA events per cell from **B**. Quantification from 10 images of 3 biological donors. Statistics were calculated using a two-tailed Student’s *t*-test. **D** Confocal images of PLA control samples which were incubated with only the secondary PLA probes. Scale bars are 40 µm.

**Figure S4.**
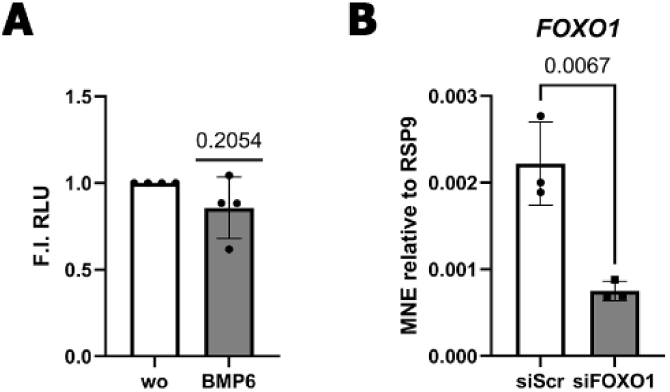
siRNA-mediated depletion of FOXO1 in human aortic endothelial cells (HAoECs) is efficient. **A** Reporter gene assay in HEK293T cells stimulated with 5 nM BMP6 for 24 h using a genomic region of the *EDN1* promoter (n=4). Values are fold-inductions over the unstimulated control. Statistics were calculated as a one-sample *t*-test. **B** qPCR analysis of FOXO1 in HAoECs exposed to oscillatory flow upon siRNA-mediated depletion of FOXO1 (n=3). Values are depicted as mean normalized expression +/- standard deviation. Statistics were calculated as a two-tailed Student’s *t*-test.

**Figure S5.**
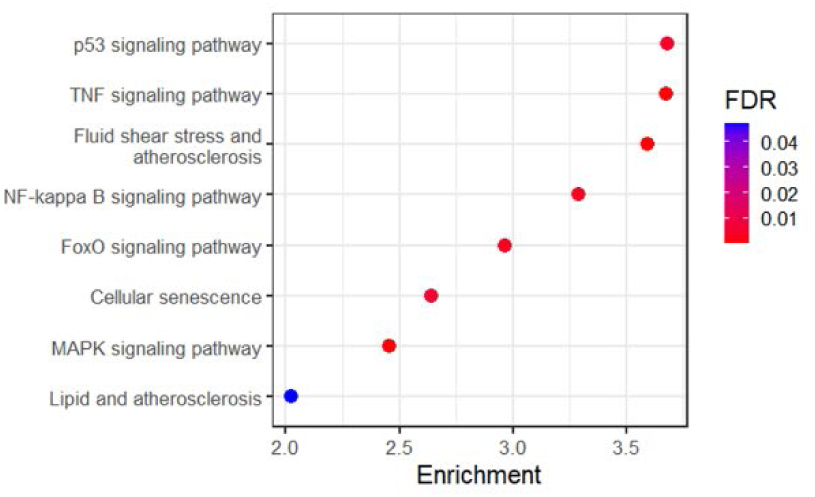
Inflammatory and atherosclerosis associated pathways depend on FOXO1 activity. Gene ontology enrichment of KEGG pathway terms determined from differentially expressed genes (DEGs) (adjusted p-value < 0.05, fold change > |log_2_(0.585)|) from RNA-Seq data of HAoECs exposed to oscillatory flow (OS) in absence or presence of FOXO1 inhibitor AS1842856 (0.5 µM).

